# Atg39 links and deforms the outer and inner nuclear membranes in selective autophagy of the nucleus

**DOI:** 10.1101/2021.03.29.437603

**Authors:** Keisuke Mochida, Toshifumi Otani, Yuto Katsumata, Hiromi Kirisako, Chika Kakuta, Tetsuya Kotani, Hitoshi Nakatogawa

## Abstract

In selective autophagy of the nucleus (hereafter nucleophagy), nucleus-derived double membrane vesicles (NDVs) are formed, sequestered within autophagosomes, and delivered to lysosomes or vacuoles for degradation. In *Saccharomyces cerevisiae*, the nuclear envelope (NE) protein Atg39 acts as a nucleophagy receptor, which interacts with Atg8 to target NDVs to forming autophagosomal membranes. In this study, we revealed that Atg39 is anchored to the outer nuclear membrane (ONM) via its transmembrane domain and also associated with the inner nuclear membrane (INM) via membrane-binding amphipathic helices (APHs) in its perinuclear space region, thereby linking these membranes. We also revealed that overaccumulation of Atg39 causes the NE to protrude towards the cytoplasm, and the tips of the protrusions are pinched off to generate NDVs. The APHs of Atg39 are crucial for Atg39 assembly in the NE and subsequent NE protrusion. These findings suggest that the nucleophagy receptor Atg39 plays pivotal roles in NE deformation during the generation of NDVs to be degraded by nucleophagy.

## Introduction

Macroautophagy (hereafter autophagy) is the mechanism whereby cellular material such as proteins and RNA as well as larger structures such as protein aggregates, phase-separated liquid droplets, and membrane-bound organelles are degraded, playing an important role in the maintenance and regulation of various cellular functions (Nakatogawa, 2020; Morishita and Mizushima, 2019). Autophagy initiates with the formation and expansion of the membrane cisterna known as the isolation membrane or phagophore, which bends, becomes spherical, and seals to sequester degradation targets within the resulting double membrane vesicle, the autophagosome. The autophagosome then fuses with the lysosome in animals and the vacuole in yeast and plants for degradation of the sequestered material within these lytic organelles. Sequestration of cellular components into autophagosomes proceeds in both a selective and non-selective manner. In selective types of autophagy (hereafter selective autophagy), autophagy receptors bind to degradation targets and act to recruit core Atg proteins, which mediate autophagosome biogenesis, to the targets for their efficient sequestration into the autophagosomes (Morishita and Mizushima, 2019; Farré and Subramani, 2016). Previous studies suggest that different organelles, including mitochondria, peroxisomes, the endoplasmic reticulum (ER), and the nucleus, are degraded by selective autophagy, and autophagy receptors responsible for degradation of these organelles have also been identified (Morishita and Mizushima, 2019; Farré and Subramani, 2016).

The nucleus contains chromosomes, and the double membraned nuclear envelope (NE) separates specific reactions such as DNA replication and gene transcription from the cytoplasm. The outer nuclear membrane (ONM) and the perinuclear space (NE lumen) are continuous with the membrane and the lumen of the ER, respectively, and therefore, these membranes and spaces share many proteins. The ubiquitin-proteasome system is vital in preventing the accumulation of aberrant proteins in the nucleus. However, nuclear inclusion bodies are formed when these proteins accumulate beyond the capacity of the system in neurodegenerative diseases such as Huntington’s disease (Enam et al., 2018; Woulfe, 2008). Recent studies suggest that nuclear components are delivered to and degraded in lysosomes or vacuoles via autophagy (Mijaljica and Devenish, 2013; Mochida et al., 2015). In the budding yeast *Saccharomyces cerevisiae*, we identified the transmembrane protein Atg39 as a receptor for selective autophagy of the nucleus (hereafter nucleophagy) (Mochida et al., 2015). Meanwhile, yeast cells lacking Atg39 exhibited abnormal nuclear morphology and an early cell death phenotype under nitrogen starvation. We and other groups also reported selective autophagy of nuclear pore complexes and nuclear lamina in yeast and mammalian cells, respectively (Dou et al., 2015; Tomioka et al., 2020; Lee et al., 2020). These results therefore suggest that selective degradation of nuclear components via autophagy also has a significant impact on nuclear homeostasis in cells.

We previously revealed that when *S. cerevisiae* cells are subjected to nitrogen starvation or treated with the Tor kinase complex 1 (TORC1) inhibitor rapamycin, Atg39 is expressed and localized to the NE, and, as with other autophagy receptors (Farré and Subramani, 2016), it interacts with Atg11, which serves as a scaffold to recruit core Atg proteins, and Atg8, which is located on expanding isolation membranes, loading nucleus-derived double membrane vesicles (NDVs) of ∼200 nm into the autophagosomes (Mochida et al., 2015). The outer and inner membranes of NDVs are derived from the ONM and inner nuclear membrane (INM), respectively, with nucleoplasmic and nucleolar proteins existing in the vesicle lumen. To generate NDVs, the ONM and INM need to induce deformation and fission in a coordinated manner. Because NDVs do not accumulate in the cytoplasm of cells deficient for autophagosome formation, these vesicles are thought to form concomitantly with autophagosome formation. We also found that Atg39 accumulates in the NE at contact with the site of autophagosome formation; however, the mechanisms underlying these processes of NDV formation during nucleophagy remain unknown.

In this study, we discovered that Atg39 binds to both the ONM and INM via its N-terminal transmembrane domain and C-terminal amphipathic helices (APHs), respectively, thereby linking these membranes. We also found that overexpression of Atg39 drives the formation of protrusions from the NE accompanied by both the ONM and INM. The APHs in the C-terminal perinuclear space region of Atg39 play multiple roles in nucleophagy: Atg39 retention to the NE; the basal and enhanced assembly of Atg39 in the NE, the latter of which occurs in conjunction with autophagosome formation; and the formation of NE protrusions, the tips of which are pinched off to generate NDVs.

## Results

### Atg39 is an ONM protein

Although Atg39 was predicted to be an integral membrane protein with a single transmembrane domain, separating it into N-terminal (1–144) and C-terminal (165–398) regions, its membrane topology has yet to be demonstrated experimentally. To determine the membrane topology of Atg39, lysates were prepared from yeast cells expressing Atg39 N- and C-terminally tagged with HA and GFP sequences, respectively, were treated with proteinase K in the presence or absence of the detergent Triton X-100 (TX-100) (Fig. 1A and B). Immunoblotting using anti-HA antibody showed that the HA tag in HA-Atg39-GFP was digested by proteinase K regardless of the presence of TX-100 (Fig. 1A). In contrast, immunoblotting with anti-GFP antibody revealed that the GFP tag in Atg39-GFP was largely resistant to proteinase K in the absence of TX-100, although it was trimmed to a size corresponding to the protein lacking the region N-terminal to the transmembrane domain (Fig. 1B). These results suggest that Atg39 is a single-pass membrane protein with N- and C-terminal regions exposed to the cytoplasm and the perinuclear space, respectively (Fig. 1C), consistent with the existence of Atg8- and Atg11-binding sequences in the N-terminal region (Mochida et al., 2015).

**Figure 1.**
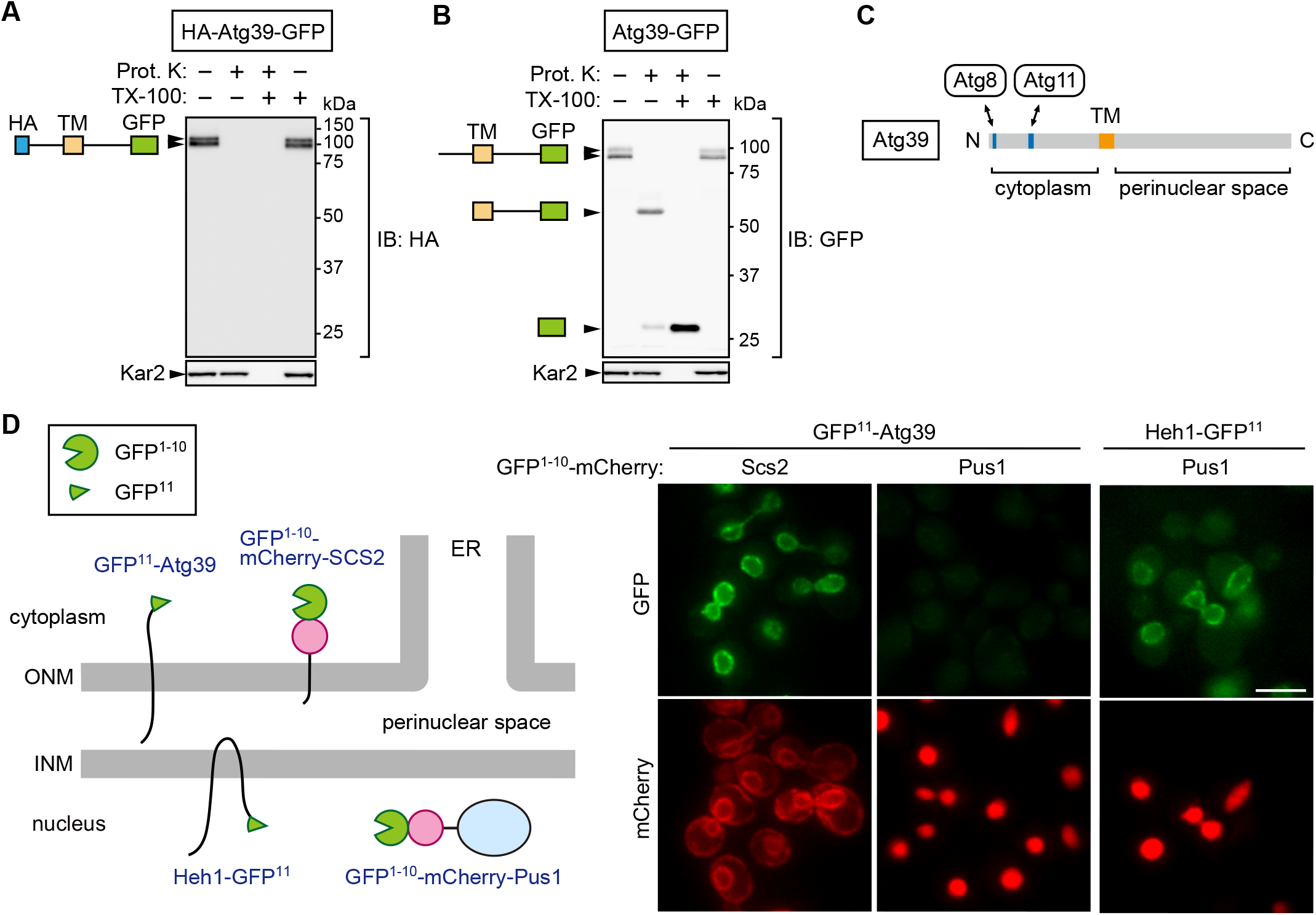
Atg39 is a transmembrane protein embedded in the ONM. **(A, B)** Proteinase K protection assay was performed to determine the membrane topology of Atg39. Lysates were prepared from cells expressing HA-Atg39-GFP (A) or Atg39-GFP (B) and incubated with proteinase K (Prot. K) in the absence or presence of Triton X-100 (TX-100). N-terminal HA and C-terminal GFP tags were detected by immunoblotting using anti-HA and anti-GFP antibodies, respectively. Kar2, an ER lumenal protein. **(C)** Schematic diagram of Atg39. TM, transmembrane domain. **(D)** Split GFP-based assay was carried out to determine the localization of Atg39. An N-terminal fragment of GFP (GFP^1-10^) and mCherry were fused to a transmembrane domain of Scs2 (GFP^1- 10^-mCherry-Scs2) or Pus1 (GFP^1-10^-mCherry-Pus1), and the remaining C-terminal fragment of GFP (GFP^11^) was fused to the N-terminus of Atg39 or the C-terminus of the INM protein Heh1. Cells expressing these proteins were observed under a fluorescence microscope. Scale bars, 5 µm.

To investigate whether Atg39 is embedded in the ONM only or in both the ONM and INM, we used a split-GFP-based system (Smoyer et al., 2016). In this system, an N-terminal sequence of GFP containing 10 β strands (GFP^1-10^) was attached to the ONM/ER membrane protein Scs2 (GFP^1-10^-mCherry-Scs2) or the nucleoplasmic protein Pus1 (GFP^1-10^-mCherry-Pus1), and the remaining C-terminal sequence containing the last β strand of GFP (GFP^11^) was fused to the N terminus of Atg39 (GFP^11^-Atg39) (Fig. 1D). When the two GFP fragments exist in the same compartment, they form a functional GFP. GFP fluorescence was observed in the NE of cells coexpressing GFP^1-10^-mCherry-Scs2 and GFP^11^-Atg39, but not those expressing GFP^1-10^- mCherry-Pus1 instead of GFP^1-10^-mCherry-Scs2. When GFP^1-10^-mCherry-Pus1 was coexpressed with GFP^11^ fused to the INM protein Heh1 (also known as Src1), GFP fluorescence was observed in the NE (INM), confirming that GFP^1-10^-mCherry-Pus1 can form a functional GFP if the GFP^11^- fused protein exists in the same compartment. These results support the conclusion that Atg39 specifically localizes to the ONM.

### APHs of Atg39 bind to the INM to retain Atg39 in the NE

Because the ONM is continuous with the ER membrane, a mechanism must exist that retains Atg39 in the ONM. The determined topology of Atg39 suggests that the C-terminal region (165-398) is exposed to the perinuclear space; however, the importance and function of this region in nucleophagy have yet to be investigated. We therefore examined the intracellular localization of C-terminally truncated Atg39 mutants (Fig. 2A, B). Whereas Atg39^1-347^ and Atg39^1-325^, as with wild-type Atg39, exhibited specific localization to the ONM, Atg39^1-296^ and Atg39^1-194^ leaked out into the ER, which spreads mainly beneath the plasma membrane in yeast cells (West et al., 2011). In contrast, deletion of the N-terminal cytoplasmic region did not affect the ONM localization of Atg39 (Fig. S1A). These results suggest that the C-terminal region 297-325 is responsible for limiting Atg39 localization to the ONM.

**Figure 2.**
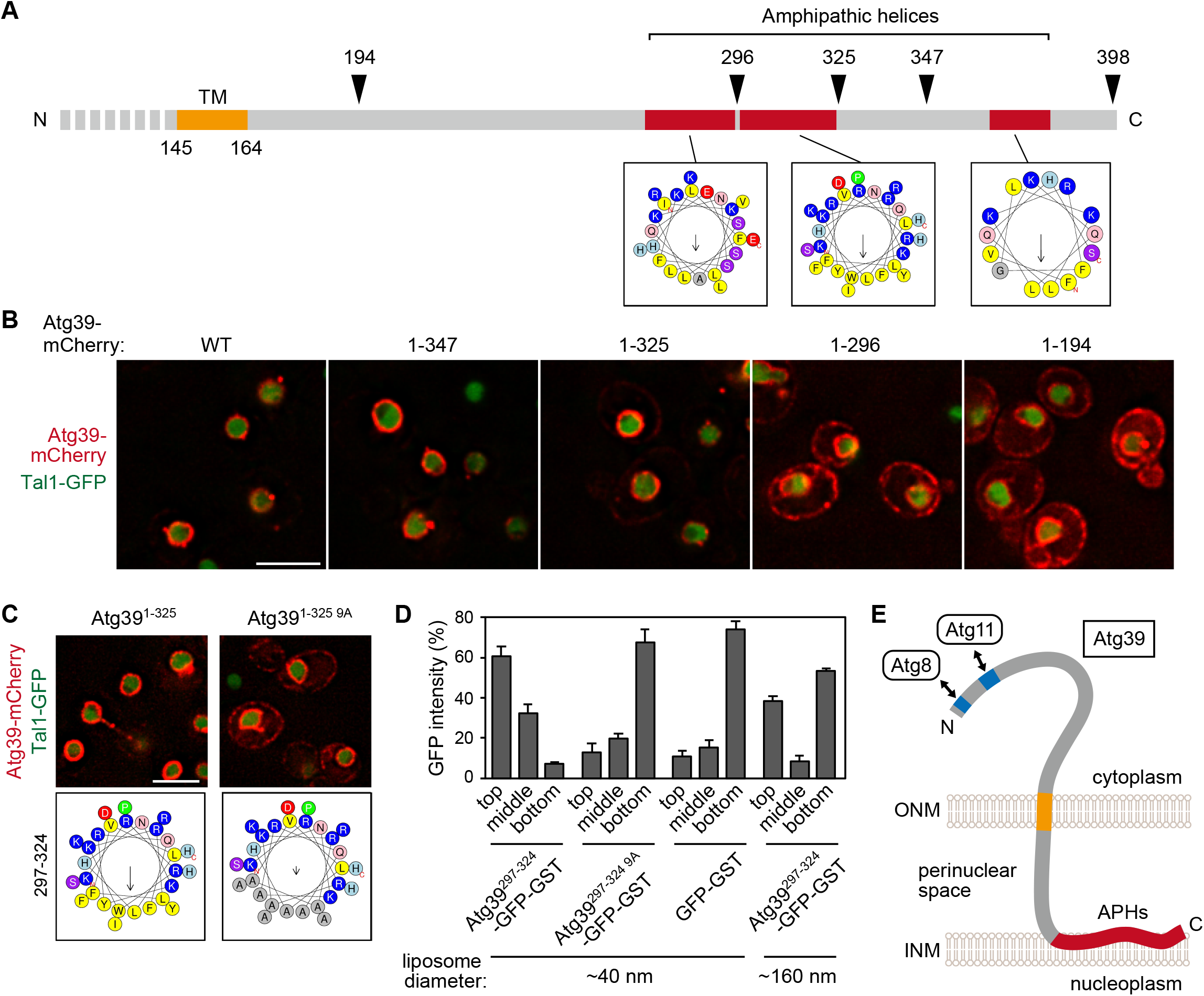
The APHs of Atg39 bind to the INM to retain Atg39 at the ONM. **(A)** Schematic diagram of the C-terminal perinuclear space region of Atg39. Helical wheels were generated using a HeliQuest server. **(B)** mCherry-fused wild-type Atg39 (WT) and C-terminally truncated mutants were expressed under the constitutive *HRR25* promoter and analyzed by fluorescence microscopy. Tal1, a nucleoplasmic protein. **(C)** The localization of Atg39^1-325^-mCherry and Atg39^1-325 9A^-mCherry was analyzed by fluorescence microscopy. **(D)** Liposome flotation assay was carried out to examine the membrane-binding ability of APH^297-325^. Purified GFP-GST-fused proteins were mixed with small (∼40 nm) or large (∼160 nm) liposomes, and then the liposomes and liposome-bound proteins were floated by ultracentrifugation. Data are shown as means ± s.d. (n = 3). **(E)** The determined membrane topology of Atg39. Scale bars, 5 µm.

In analysis of the localization of N-terminally truncated Atg39 mutants, we found that the mutant lacking cytoplasmic and transmembrane domains (Atg39^167-398^) localized to the surface of lipid droplets (Fig. S1A, B). Although this is an artifact of cytoplasmic expression of the perinuclear space region, it raised the possibility that the perinuclear space region of Atg39 has the ability to associate with lipid membranes. The C-terminal region of Atg39 is predicted to largely form a helical conformation, containing at least three APHs, including helix 297-324 (APH^297-324^), which is important for the ONM localization of Atg39 (Fig. 2A, B). APHs are known to bind to membranes through their hydrophobic surface (Drin and Antonny, 2010). We therefore speculated that APH^297-324^ associates with the INM from the side of the perinuclear space to prevent Atg39 from leaking into the ER. To examine this possibility, nine hydrophobic residues in APH^297-324^ were replaced with alanine (9A) (Fig. 2C). Similar to Atg39^1-296^, this 9A mutant mislocalized to the ER, highlighting the importance of the amphipathic property of APH^297-324^ in Atg39 retention to the ONM.

Next, we performed a co-flotation assay using liposomes and purified APH^297-324^ tagged with GFP-GST (Fig. 2D). Although APH^297-324^-GFP-GST co-floated with small liposomes (∼40 nm in diameter) after centrifugation, the 9A mutation severely impaired liposome binding of this fusion protein, demonstrating that APH^297-324^ indeed has a membrane-binding ability. To clarify that the APHs of Atg39 associate with the INM in the perinuclear space, we performed immunoelectron microscopy of an Atg39 variant in which the HA sequence was inserted into the region between APH^297-324^ and APH^365-379^ (Fig. S1C). Reaction of the immunogold particles with the HA sequence was frequently detected in the vicinity of the INM. Overall, these findings suggest that, in addition to the transmembrane domain, which penetrates the ONM, Atg39 contains membrane-binding APHs and is exclusively localized to the ONM via the insertion of APHs into the perinuclear space-facing layer of the INM (Fig. 2E). These two types of membrane-binding domains bridging the ONM and INM allowed us to speculate that Atg39 couples deformation of the ONM and INM during NDV formation in nucleophagy.

### APHs of Atg39 promote Atg39 assembly at the NDV formation site

Next, we examined nucleophagy in APH-deleted Atg39 mutants. Nucleophagy activity can be evaluated by the amounts of protease-resistant GFP fragments (GFP’) generated by vacuolar degradation of the nucleoplasmic protein Tal1 (Breker et al., 2014) fused with GFP. The results showed that the Atg39^1-296^ and Atg39^1-194^ mutants were severely defective in nucleophagy and that the Atg39^1-347^ mutant had a milder defect (Fig. 3A), confirming that the APHs of Atg39 are important for nucleophagy.

**Figure 3.**
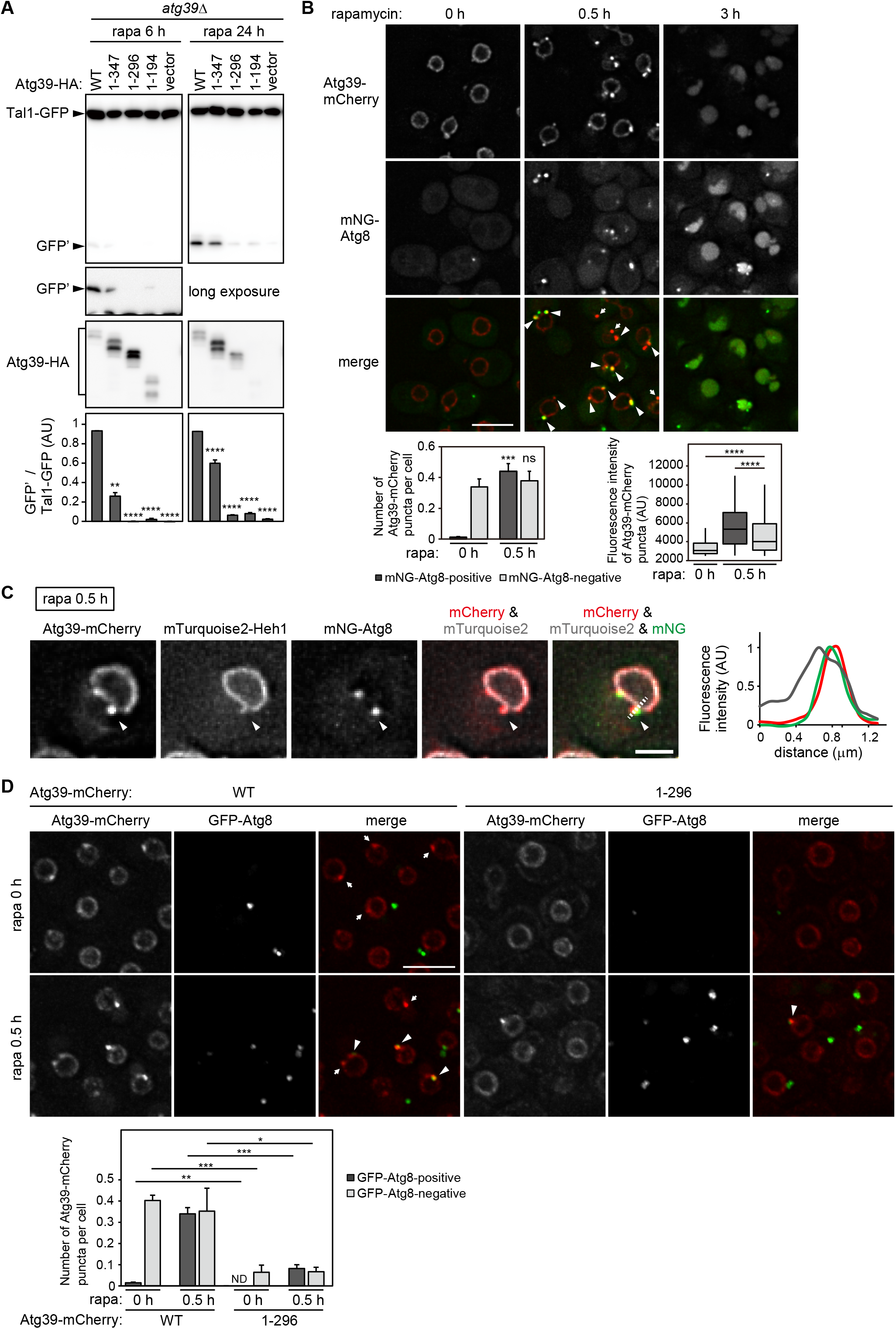
The APHs of Atg39 promote Atg39 assembly at the site of NDV formation. **(A)** Nucleophagy activity in *atg39*Δ cells expressing wild-type (WT) Atg39 tagged with the HA sequence or C-terminally truncated mutants. Cells were treated with rapamycin to induce nucleophagy, and degradation of Tal1-GFP was examined by immunoblotting using an anti-GFP antibody. The quantification results are shown as means ± s.d. (n = 3). \*\*\*\**P* < 0.0001; \*\**P* < 0.01 (Student’s *t*-test). **(B)** Cells constitutively expressing Atg39-mCherry were treated with rapamycin and observed under a fluorescence microscope. The number of mNG-Atg8-positive and negative puncta of Atg39-mCherry (bottom left), and the fluorescence intensity of these puncta (bottom right) were measured and are shown graphically. Bars represent means ± s.d. (n = 3) (bottom left). \*\*\**P* < 0.001 (Student’s *t*-test) (bottom left). \*\*\*\**P* < 0.001 (Mann–Whitney *U* test) (bottom right). **(C, D)** Cells constitutively expressing Atg39-mCherry were treated with rapamycin for 0.5 h and analyzed by fluorescence microscopy. Fluorescence intensity along the dashed line in (C) was measured and is graphically shown. The bars represent means ± s.d. (n = 3). \**P* < 0.05; \*\**P* < 0.01; \*\*\**P* < 0.001 (Student’s *t*-test). Arrow heads, mNG/GFP-Atg8-positive Atg39-mCherry puncta. Arrows, mNG/GFP-Atg8-negative Atg39 puncta. Scale bars, 5 µm (B, D), 2 µm (C).

Next, we determined which nucleophagy step is defective in Atg39 mutants lacking the APHs. To this end, we first determined the intracellular dynamics of Atg39 during the process of nucleophagy. Because the expression of Atg39 is repressed under normal conditions (Mochida et al., 2015), in these experiments, we expressed Atg39 using the constitutive *HRR25* promoter and observed changes in Atg39 localization upon rapamycin treatment (nucleophagy induction). Before adding rapamycin, Atg39-mCherry was almost uniformly distributed in the NE, although a significant proportion of cells had faint Atg39-mCherry puncta that did not colocalize with the autophagosomal membrane marker Atg8 tagged with mNeonGreen (mNG) (Fig. 3B). In contrast, 30 min after rapamycin addition, Atg39-mCherry assembled and formed bright puncta that colocalized with mNG-Atg8, and after 3 h, these puncta disappeared, and mCherry and mNG fluorescence was observed within the vacuole, showing the progression of nucleophagy (Fig. 3B). Moreover, after 30 min of rapamycin treatment, the NE frequently formed protrusions (or buds), with notable accumulation of Atg39-mCherry (Fig. 3B). Colocalization of the INM protein Heh1 suggests that the INM protrudes together with the ONM (Fig. 3C). These results further suggest that NE protrusions associated with bright Atg39-mCherry puncta represent forming NDVs. In contrast, the bright puncta of Atg39-mCherry did not form in cells lacking the core Atg protein Atg1 or Atg2, and this was also the case in cells lacking the adapter protein Atg11, which recruits the core Atg proteins (Fig. S2). Meanwhile, the formation of Atg8-negative, faint Atg39-mCherry puncta, which did not increase following rapamycin treatment (Fig. 3B), was largely unaffected by deletion of genes encoding these proteins (Fig. S2). These results suggest that Atg39 partially assembles in the NE, and that this assembly is strongly enhanced in line with the formation of the autophagosomal membrane.

As with the membrane-binding APHs analyzed previously (Drin and Antonny, 2010), APH^297-324^ of Atg39 preferentially bound to smaller liposomes (membranes with high positive curvature) (Fig. 2D). Because protruding membrane domains have higher curvature than other parts of the NE, it is possible that the APHs of Atg39 sense the local curvature in the INM during assembly. Indeed, deletion of the APHs decreased the formation of both faint (rapamycin-independent, Atg8-negative) and bright (rapamycin-dependent, Atg8-positive) Atg39-mCherry puncta (Fig. 3D). These results suggest that the APHs of Atg39 are involved in both basal and enhanced Atg39 assembly in the NE. As mentioned above, enhanced Atg39 assembly depends on autophagosome formation (Fig. S2), and thus it is likely that autophagosome formation in the cytoplasm and APHs bound to the INM act together to assemble Atg39 at the site of NDV formation.

### APHs of Atg39 are involved in deformation of the NE during NDV formation

The insertion of APHs into one side of the lipid bilayer can bend the membrane (Drin and Antonny, 2010). We therefore examined whether Atg39 itself can induce local NE remodeling for the generation of NDVs. Atg39-mCherry was overexpressed using the copper-inducible *CUP1* promoter in *atg1*Δ cells to block nucleophagy and accumulate Atg39-mCherry in the NE. We found that the accumulation of Atg39 in the NE caused the extension of tubules from the NE (Fig. 4A), consistent with a previous report (Vevea et al., 2015). Rapamycin treatment increased the formation of these NE tubules in terms of both number and length (Fig. 4A) in *atg1*Δ cells. In addition, deletion of neither *ATG8* nor *ATG11* affected this tubule formation (Fig. S3A), suggesting that rapamycin treatment promotes NE tubule formation independent of autophagosome formation and the interactions of Atg39 with Atg8 and Atg11. Electron microscopy further confirmed that both the ONM and INM join to form these tubules (Fig. 4B). The tips of the tubules were often swollen and contained components similar to the nucleoplasm, consistent with the idea that these structures are related to NDV formation. We also showed that the formation of NE tubules was much less efficient in cells overexpressing Atg39^1-296^ compared with the wild-type Atg39, suggesting that the APHs of Atg39 are involved in NE tubulation (Fig. 4C). Taken together, these findings suggest that Atg39 has the ability to induce the protrusion of both the ONM and INM to form NDVs, with the perinuclear-space APHs playing an important role. Although NE tubulation is an exaggerated phenomenon caused by Atg39 overexpression, puncta of endogenous Atg39 were often observed at the tip of the NE protrusions (Fig. S3B). Furthermore, time-lapse microscopy showed that budding and fission of Atg39-mCherry-enriched structures occur at the tip of NE tubules (Fig. S3C). Overall, these results suggest that endogenous Atg39 also deforms the NE and generates short protrusions, the tip of which provides a site for NDV formation.

**Figure 4.**
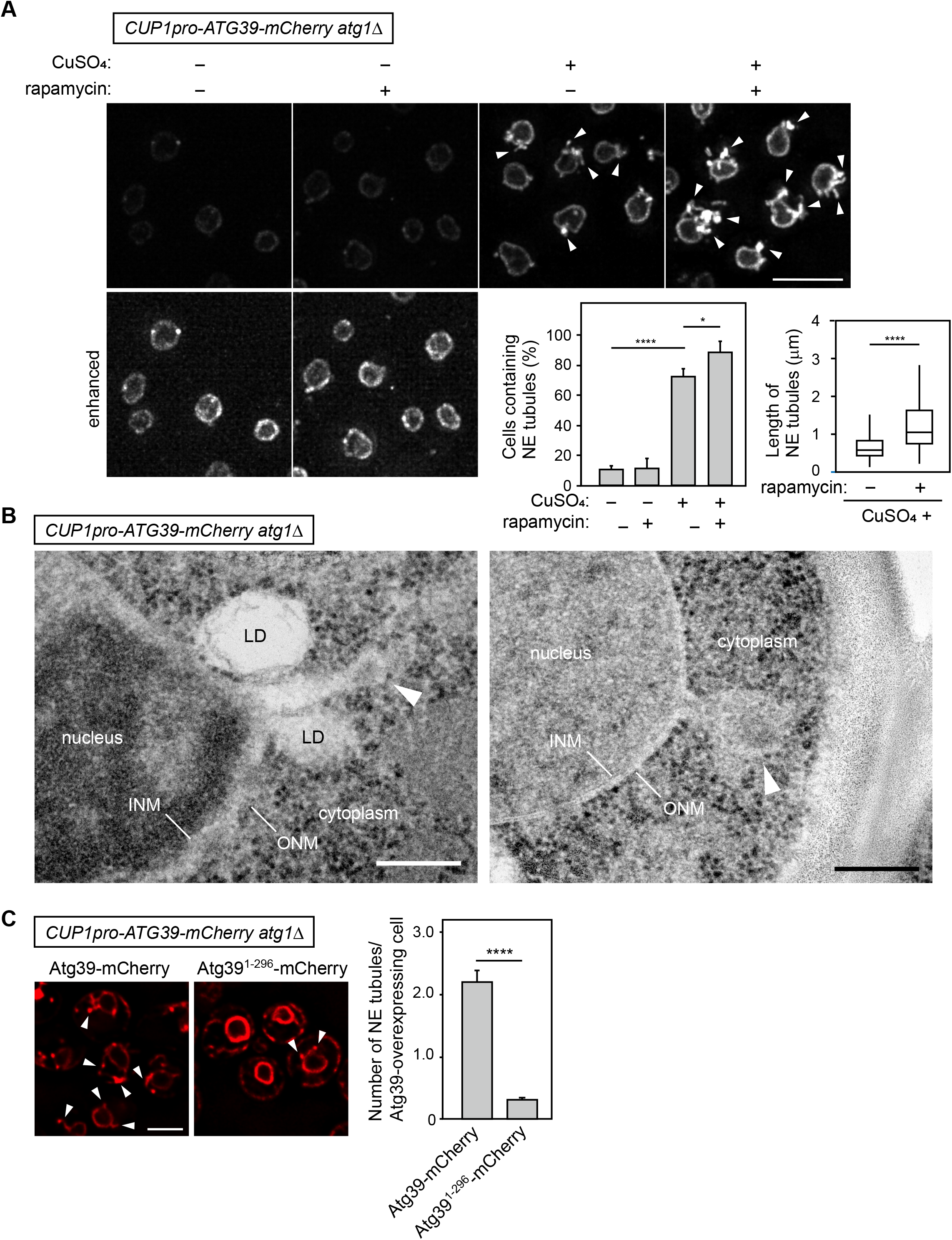
Atg39 accumulation in the NE causes NE tubulation dependent on the APHs. **(A)** *atg1*Δ cells expressing Atg39-mCherry under the *CUP1* promoter were grown in the presence or absence of 250 µM CuSO_4_ overnight and treated with rapamycin for 2 h. The percentage of cells containing NE tubules (left) and length of NE tubules (right) are shown. Bars represent means ± s.d. (n =3). \**P* < 0.05; \*\*\*\**P* < 0.0001 (Tukey’s multiple comparisons test) (left). \*\*\*\**P* < 0.0001 (Mann–Whitney *U* test) (right). **(B)** *atg1*Δ cells overexpressing Atg39-mCherry in the presence of copper ions were treated with rapamycin for 2 h, and the nuclear morphology of these cells was examined by electron microscopy. LD, lipid droplet. **(C)** mCherry-fused Atg39 and Atg39^1-296^ were overexpressed in *atg1*Δ cells. These cells were treated with rapamycin for 2 h, and the formation of NE tubules was analyzed by fluorescence microscopy. Arrowheads, NE tubules. \*\*\*\**P* < 0.0001 (Tukey’s multiple comparisons test). Scale bars, 5 µm (A, C), 200 nm (B).

### Chromosomes are excluded from NE tubules and NDVs

Lastly, we investigated the incorporation of nuclear components into NE tubules under Atg39 overexpression. Consistent with the electron microscopic observations showing that these tubules are composed of both the ONM and INM (Fig. 4B), fluorescence microscopy showed signals of the INM protein Heh1 in most of tubules positive for Atg39 (Fig. 5A). Moreover, although less but significant proportions of NE tubules also contained the nucleoplasmic protein Tal1 and the nucleolar protein Nop56, the histone H2A Hta2 seldom colocalized with these tubules. We also confirmed that histone proteins were absent from NDVs formed using cells lacking Ypt7, a Rab GTPase essential for autophagosome-vacuole fusion (Kirisako et al., 1999). When these cells were treated with rapamycin, numerous Atg39-mCherry puncta representing NDVs sequestered within the autophagosomes accumulated in the cytoplasm (Fig. 5B). Whereas puncta of Tal1-GFP were observed outside the nucleus, colocalizing with cytoplasmic Atg39-mCherry puncta, the GFP-fused histones Hta2 and Htz1 did not form any puncta (Fig. 5B). Moreover, Hoechst-stained DNA was not detected in Atg39-mCherry puncta in these cells (Fig. S4), suggesting that chromosomes do not enter NE protrusions and are excluded from sequestration into NDVs.

**Figure 5.**
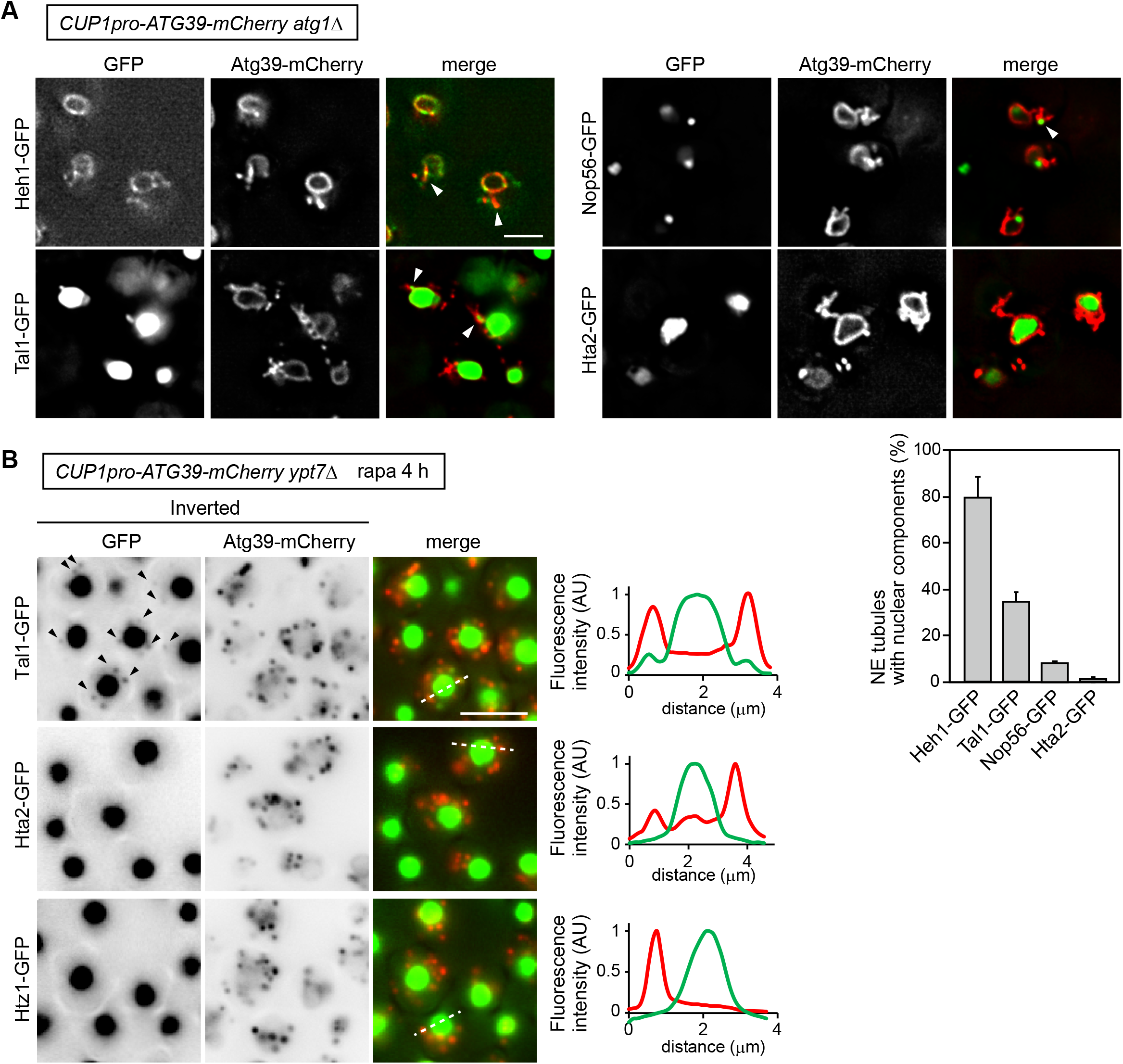
NE tubules and NDVs exclude chromosomes. **(A)** NE tubule formation was induced by overexpression of Atg39-mCherry in *atg1*Δ cells followed by treatment with rapamycin treatment for 2 h. The colocalization of Atg39-mCherry-positive NE tubules with GFP-fused Heh1, Tal1, Nop56, and Hta2 was examined by fluorescence microscopy. Arrowheads, NE tubules containing GFP signals. The percentage of NE tubules containing nuclear components is shown in the graph. Bars represent means ± s.d. (n =3). **(B)** *ypt7*Δ cells overexpressing Atg39-mCherry were treated with rapamycin for 4 h. Arrowheads, puncta of Tal1-GFP that colocalized with those of Atg39-mCherry. Fluorescence intensity along the dashed line was measured, and the results are shown in the graph. Scale bars, 5 µm

## Discussion

In this study, we showed that the nucleophagy receptor Atg39 is a single membrane-spanning ONM protein whose N-terminal cytoplasmic region interacts with Atg8 and Atg11 and whose C-terminal perinuclear space region contains APHs that associate with the INM. Deletion of these APHs resulted in Atg39 mislocalization to the ER, with impaired basal (autophagosome formation-independent) and enhanced (autophagosome formation-dependent) assembly in the NE as well as reduced nucleophagy activity. Overexpression of Atg39 induced the extension of double membrane tubules from the NE depending on the APHs. Based on these results, we propose the following model for the mechanism of NDV formation during Atg39-mediated nucleophagy (Fig. 6). First, a certain amount of Atg39 assembles in the NE in a manner dependent on its APHs and independent of autophagosome formation (GFP-Atg8-negative Atg39-mCherry puncta). Dependency of this basal Atg39 assembly on the APHs suggests that at least the INM protrudes into the perinuclear space at this stage. The APHs of Atg39 and/or other proteins (see below) may also be involved in this initial deformation of the NE/INM. The cytoplasmic region of Atg39 in the assemblage subsequently recruits the core Atg proteins via Atg11, thereby initiating autophagosome formation on the NE. Atg39 then assembles further, probably via interaction with Atg11 and the core Atg proteins associated with the isolation membrane such as Atg8, thereby inducing protrusion of the INM into the perinuclear space through the APHs while remaining anchored to the ONM in the transmembrane domain, resulting in the formation of double membrane protrusions out of the NE. Both the ONM and INM cause fission at the tip of the protrusions, resulting in the formation of NDVs containing intranuclear components, which are then sequestered into the autophagosomes and delivered to the vacuole for degradation.

**Figure 6.**
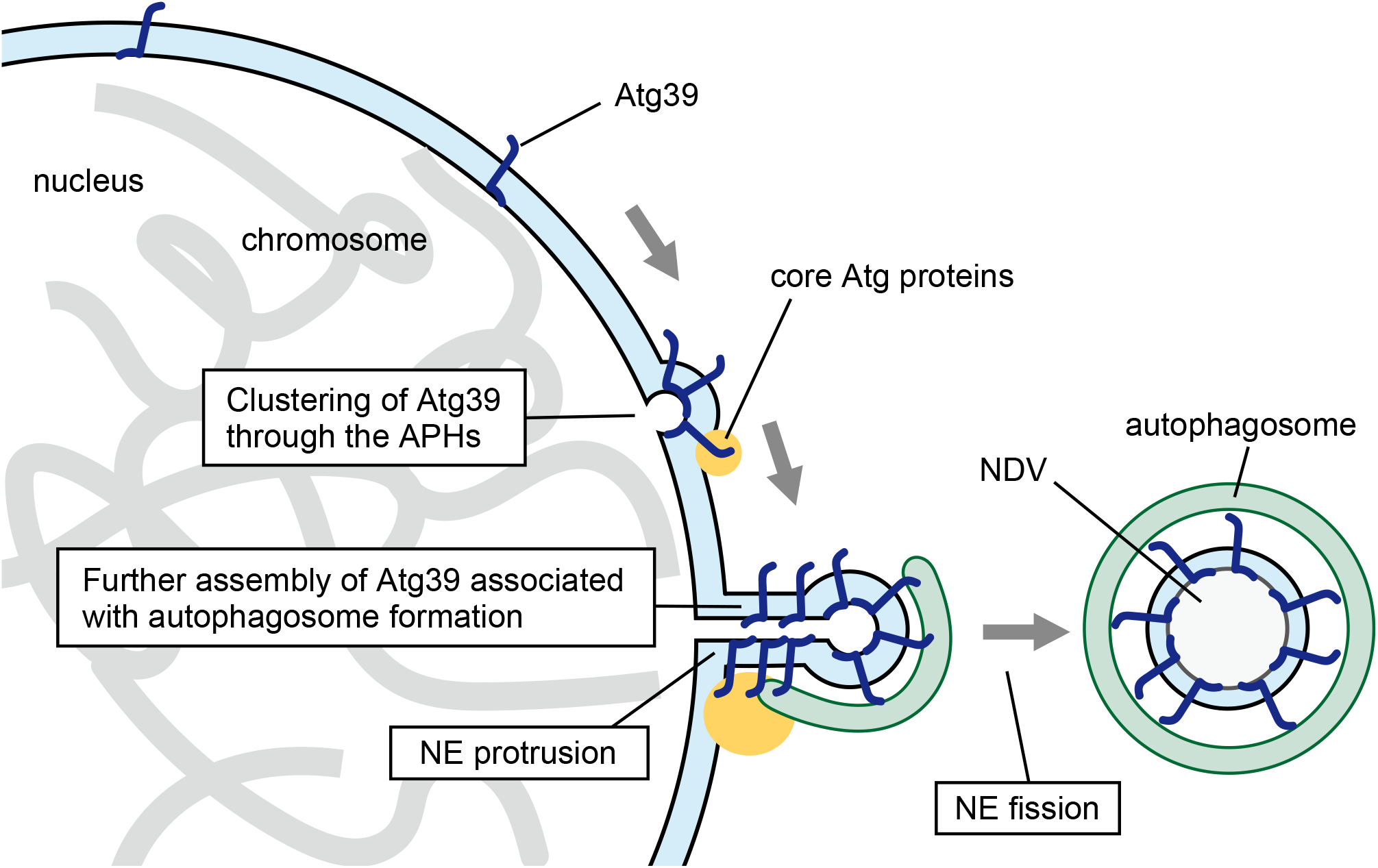
Model mechanism of NDV formation in Atg39-mediated nucleophagy. Atg39 forms a small cluster depending on APHs in the NE region where the INM partially protrudes via an unknown mechanism. Then, Atg39 in the cluster recruits the core Atg proteins via Atg11 on the cytoplasmic surface of the NE, initiating autophagosome formation, which triggers the further assembly of Atg39, probably via interaction with Atg11 and Atg8 on the forming autophagosomal membrane. The condensation of Atg39 locally enhances its APH-dependent membrane-deforming activity, protruding the NE toward the cytoplasm at the site. An unknown mechanism mediates fission of the tip of the protrusion to release a NDV, which is sequestered into the autophagosome via the interaction between Atg39 and Atg8, a canonical function of autophagy receptors.

Although membrane bending via the wedging effect of APH insertion has been well documented, the findings suggest that the perinuclear space region of Atg39 also generates membrane curvature through an additional mechanism that can cooperate with the wedging mechanism. For example, a number of BAR proteins contain APHs and form crescent-shaped dimers, which serve as scaffolds and act together with the APHs to tubulate membranes (Baumgart et al., 2011). Steric pressure generated by local crowding of peripheral membrane proteins can also cause the tubulation of flat membranes (Stachowiak et al., 2012). Structural and in vitro studies of the perinuclear space region of Atg39 should further clarify the detailed mechanisms by which this region drives NE deformation. In addition, it is also possible that Atg39 in the cytoplasm and perinuclear space recruits other membrane-deforming proteins to generate NDVs. It should be noted that tubules extending without the INM were also observed in Atg39-overexpressing cells (Fig. 5A, S3D). These ONM tubules were not formed when Atg39 lacking the APHs was overexpressed. This observation may be merely an artifact of Atg39 overexpression but imply that the APHs of Atg39 are also involved in ONM deformation. However, because the perinuclear space surface of the protruding ONM has negative curvature, the perinuclear space region other than the APHs might contribute to deformation of the ONM.

Although the mechanism of NDV fission from the NE remains unknown, the ESCRT-III complex is a potential candidate for fission of the INM. ESCRT-III proteins form spiral filaments and mediate the invagination and fission of intralumenal vesicles in late endosomes, and these proteins are also known to exist on the nucleoplasmic surface of the INM, where they cause NE deformation similar to that observed in nucleophagy (McCullough et al., 2018; Lee et al., 2012; Webster et al., 2014). During the nucleus-to-cytoplasm exit of some herpesviruses, the INM, in association with nucleocapsids, invaginates and causes fission depending on ESCRT-III to release nucleocapsid-containing vesicles into the perinuclear space (Arii et al., 2018; Lee et al., 2012). Similarly, ESCRT-III may contribute to initial deformation, protrusion, and fission of the INM during NDV formation in nucleophagy. The molecular mechanism of ONM fission also remains unclear, but is thought to involve membrane-deforming proteins such as dynamin-related proteins and reticulon-family proteins, which are reported to mediate organelle fission during selective autophagy of mitochondria and the ER, respectively (Mao et al., 2013; Khaminets et al., 2015; Mochida et al., 2020).

In this study, INM insertion of the APHs was found to be important for the NE retention of Atg39. However, continuity between the ONM and the ER membrane poses the further question of why the APHs of Atg39 associate with the INM in the NE, but not with the opposite membrane in the ER. The difference in thickness (the distance between the two opposing membranes) might be one answer to this question, given that the ER is thicker than the NE in *S. cerevisiae* (West et al., 2011), which might prevent the APHs of Atg39 from reaching the opposite membrane in the ER. Atg39 that has been synthesized in the ER is diffused to the NE and anchored to the INM via the APHs or degraded in the ER due to unstable APHs without membrane insertion. It is also conceivable that Atg39 APHs disfavor the lumen-facing leaflet of the ER due to its flatness or negative curvature, especially with the tubular ER. The perinuclear space region of Atg39 may also contain a sequence interacting with an INM protein, which cooperates with the APHs in Atg39 anchoring to the INM.

Although nucleophagy is thought to be advantageous for cells in that it can serve as a powerful system for quality and quantity control of nuclear components, it must avoid degrading components such as chromosomes that are essential for cell viability. In this study, we showed that NE protrusions induced by overexpression of Atg39, as well as NDVs, do not contain histones and DNA. We speculate that NE protrusions are thin enough to block the entry of chromosomes into NDVs, guaranteeing the safety in degradation of the nucleus.

NDV formation during nucleophagy is associated with complicated membrane dynamics in the NE. This study focused on the APHs of Atg39, providing a deeper understanding of the underlying mechanism. However, unresolved issues remain, including how NE deformation and autophagosome formation cooperate as well as how membrane fission occurs to release NDVs from NE protrusions. Further analysis of Atg39 and identification of other proteins involved in nucleophagy will be important in addressing these issues.

## Materials and methods

### Yeast strains and plasmids

Yeast strains used in this study are listed in Table S1. Gene deletion and tagging were based on a standard PCR-based method (Janke et al., 2004). pRS303-derived plasmids were integrated into the genome at the *HIS3* locus after linearization using the restriction enzyme *Nsi*I (Sikorski and Hieter, 1989; Janke et al., 2004). Plasmids used in this study are listed in Table S2. They were constructed by amplifying appropriate DNA fragments by PCR and assembling them using the Gibson Assembly method (New England Biolabs). pRS315-NOP1pro-GFP^1-10^-mCherry-PUS1 and pRS315-NOP1pro-GFP^1-10^-mCherry-SCS2TM were generated using PCR products amplified from Addgene plasmids #86413 and #86419 and Addgene plasmids #86416 and #86419, respectively.

### Yeast cell growth conditions

Yeast cells were grown at 30°C in YPD medium (1% yeast extract, 2% peptone, and 2% glucose) or SD+CA medium (0.17% yeast nitrogen base without amino acids and ammonium sulphate, 0.5% ammonium sulfate, 0.5% casamino acids, and 2% glucose) appropriately supplemented with 0.002% adenine, 0.002% uracil, and 0.002% tryptophan. For nucleophagy induction, cells were grown to mid-log phase and treated with 200 ng/mL rapamycin. For Atg39 overexpression, cells in which *ATG39* was put under the control of the *CUP1* promoter were grown in SD+CA medium containing 250 µM CuSO_4_ overnight. In Fig. 3D, cells expressing mCherry-tagged wild-type Atg39 were grown in SD+CA medium containing 10 µM CuSO_4_ overnight. To develop large lipid droplets (Fig. S1B), cells were grown in SO+CA medium (0.17% yeast nitrogen base without amino acids and ammonium sulphate, 0.5% ammonium sulfate, 0.12% oleate, 0.2% Tween 40, 0.1% glucose, 1% casamino acids, 0.1% yeast extract, 0.002% adenine sulfate, 0.002% tryptophan, and 0.002% uracil) for 9 h.

### Cell lysate extraction and immunoblotting

Frozen yeast cells were resuspended in 200 mM NaOH and left on ice for 5 min. After centrifugation at 3,000 *g* and removal of the supernatants, the cell pellets were resuspended in urea SDS sample buffer [100 mM MOPS-KOH (pH 6.8), 4% SDS, 100 mM dithiothreitol (DTT), and 8 M urea] and incubated at 65°C for 10 min. Proteins were resolved by SDS-PAGE, transferred to PVDF membranes, and incubated with primary antibodies against GFP (Clontech, 632381), HA (Roche, 11867431001), mRFP (a gift from Dr. Toshiya Endo), and Kar2 (a gift from Dr. Toshiya Endo). After incubation with HRP-conjugated secondary antibodies, the blots were visualized using ImageQuant LAS 4000 (GE Healthcare).

### Fluorescence microscopy

Yeast cells were analyzed using two different fluorescence microscopy systems, as described previously (Mochida et al., 2020). The images in Fig. 1D, 5B, S1A, S3C, and S4 were acquired using an inverted microscope (IX81; Olympus) equipped with an electron-multiplying CCD camera (ImagEM C9100-13; Hamamatsu Photonics), a 150× objective lens (UAPON 150XOTIRF, NA/1.45; Olympus), a Z drift compensator (IX3-ZDC2; Olympus), and appropriate lasers and filters. For time-lapse imaging, cells were grown in the glass-bottom dish and kept at 30°C using a stage top incubator (TOKAI HIT). The images in Fig. S3C were deconvoluted by AutoQuant X3 software (Media Cybernetics). All other fluorescence microscopy images were acquired using a Delta Vision Elite microscope system (GE Healthcare) equipped with a scientific CMOS camera (pco.edge 5.5; PCO AG) and a 60× objective lens (PLAPON, NA/1.42; Olympus). Images acquired by a Delta Vision were deconvoluted using SoftWoRx software. All acquired images were analyzed using Fiji software (Schindelin et al., 2012).

### Quantification of fluorescence microscopy images

Atg39-mCherry puncta and mNG-Atg8 puncta were detected using the Find Maxima function of Fiji as described previously (Mochida et al., 2020). When the maxima of Atg39-mCherry puncta exists within 272 nm of those of GFP- or mNG-Atg8 puncta, they were classified as GFP/mNG-Atg8-positive. The number of the Atg39-mCherry-positive tubules extending from the NE was manually counted. The length of the NE tubules was measured using the segmented line selection tool of Fiji. In branched tubules, the length of the longest tubule was measured.

### Proteinase K protection assay

Yeast cells grown to mid-log phase were washed with 100 mM Tris-HCl (pH 8.0) containing 10 mM DTT and then converted to spheroplasts by incubation in 0.5×YPD containing 1 M sorbitol and 200 µg/mL zymolyase 100T (Nacalai Tesque) at 30°C for 30 min. Cells were washed with 20 mM HEPES-KOH (pH 7.2) containing 1.2M sorbitol and incubated in 0.5×YPD containing 1 M sorbitol and 200 ng/mL rapamycin for 3 h. After centrifugation, pelleted cells were resuspended in HSE buffer (20 mM HEPES-KOH (pH 7.2), 1M sorbitol, and 1mM EDTA) containing 0.5 × Complete protease inhibitor cocktail (PIC) (Roche) and were passed through a polycarbonate membrane filter with a 3-µm pore size (Merck Millipore). The supernatants (lysates) were obtained by removing cell debris by centrifugation then treated with 1% Triton X-100 and 100 µg/mL proteinase K on ice for 30 min. Proteolysis was stopped by the addition of 10 mM phenylmethylsulfonyl fluoride (PMSF), and after trichloroacetic acid precipitation, the proteins were solubilized in urea SDS sample buffer and analyzed by immunoblotting.

### Protein purification

BJ3505 cells overexpressing Atg39^297-324^-GFP-GST, Atg39^297-324 9A^-GFP-GST, or GFP-GST were grown to late-log phase and harvested. Frozen cells were resuspended in buffer A [50 mM Tris-HCl (pH 7.5), 500 mM NaCl, 1 mM EDTA, and 10% glycerol] containing 2 × PIC, 2 mM PMSF, and 1 mM DTT, disrupted using a Multi-beads Shocker (Yasui Kikai) and 0.5-mm YZB zirconia beads, and solubilized with 1% Triton X-100. The lysates were cleared by centrifugation at 100,000 *g* at 4°C for 30 min and then mixed with Glutathione Sepharose 4B (GE Healthcare). The Sepharose beads were washed with buffer B [50 mM Tris-HCl (pH 8.0), 500 mM NaCl, 1 mM EDTA, and 10% glycerol] containing 0.01% Triton X-100, and then the bound proteins were eluted with buffer B containing 0.01% Triton X-100 and 10 mM reduced glutathione.

### Liposome flotation assay

Liposomes were prepared as follows. Lipids in chloroform were mixed (55:25:20:0.005 mole percent POPC:DOPE:bovine liver PI:Rhodamine-DOPE) and dried in a glass tube. The lipid film was hydrated with buffer C [20 mM HEPES-KOH (pH 7.2) and 150 mM NaCl] at a lipid concentration of 1 mM before the suspension was repeatedly frozen and thawed. To prepare larger liposomes, the liposome suspension was extruded through a polycarbonate membrane filter with a 200-nm pore size (Merck Millipore). Smaller liposomes were prepared by sonicating the liposome suspension. The size of the liposomes was measured using a Zetasizer Nano S (Malvern Instruments).

40 µL of the prepared liposomes was mixed with 5 µL of 250 nM protein solution and 80 µL of buffer C, and incubated for 1 h at 30°C. 125 µl of 100% OptiPrep (Abbott Diagnostics Technologies AS) was added to the mixture, which was then overlaid with 450 µL of buffer C containing 40% OptiPrep and 200 µL of Optiprep-free buffer C. After centrifugation at 200,000 *g* at 4°C for 105 min, the top, middle, and bottom fractions (300 µL each) were collected, and GFP fluorescence in each fraction was measured using a Varioskan Flash (Thermo Scientific).

### Electron microscopy

Electron microscopy was performed by Tokai-EMA Inc. Yeast cells were sandwiched with copper disks, rapidly frozen at −175°C, and freeze-substituted with ethanol containing 2% glutaraldehyde and 0.5% tannic acid. After dehydration, the samples were embedded in Quetol-651 resin and ultra-thin sectioned for observation under a transmission electron microscopy (JEM-1400Plus; JEOL). For immunoelectron microscopy, the samples were prepared in a similar manner, except that cells were freeze-substituted with ethanol containing 1% tannic acid and 2% water and embedded in LR white resin. The sections were incubated with an antibody against HA (11867431001, Roche) and then with a secondary antibody-conjugated to 10-nm gold particles.

## Acknowledgements

We thank the members of our laboratory for materials, discussions, and technical and secretarial support; and the Biomaterials Analysis Division of the Open Facility Center at the Tokyo Institute of Technology for DNA sequencing. This work was supported in part by KAKENHI Grants-in-Aid for Scientific Research JP17H01430 and JP19H05708 (to H.N.) from the Ministry of Education, Culture, Sports, Science, and Technology of Japan; AMED Grant Number JP21gm1410004 (to H.N.); STAR Grant funded by the Tokyo Tech Fund (to H.N.).

The authors declare no competing financial interests.

## Author contributions

K. Mochida, T. Kotani, and H. Nakatogawa designed the project. K. Mochida, T. Otani, and Y. Katsumata performed the experiments with the help of H. Kirisako and C. Kakuta. K. Mochida and H. Nakatogawa wrote the manuscript. All authors analyzed and discussed the results and commented on the manuscript.

## Supplemental Figure Legends

**Figure S1.**
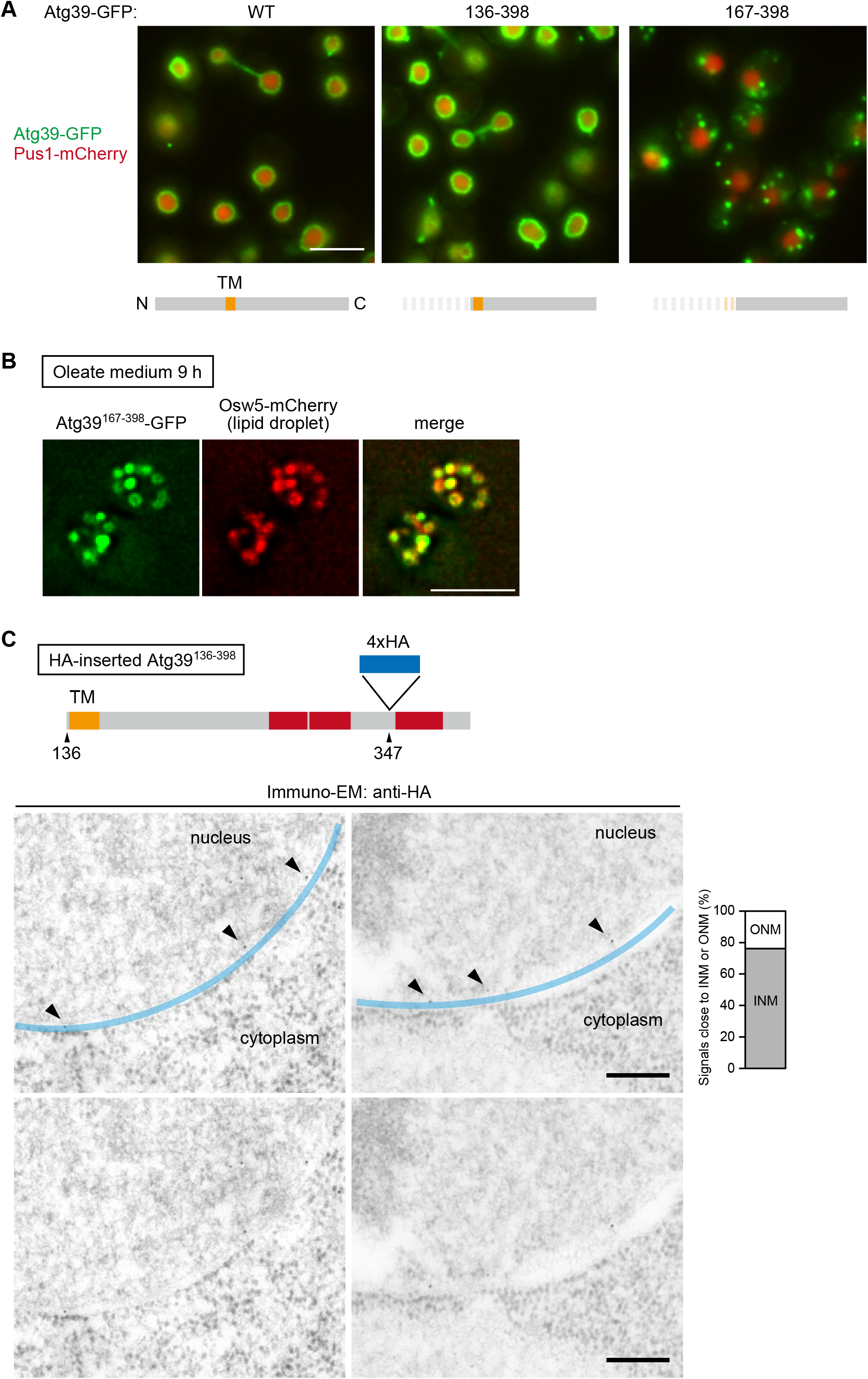
The perinuclear space region of Atg39 associates with the INM. **(A)** The intracellular localization of mCherry-fused wild-type Atg39 (WT) and the N-terminally-truncated mutants expressed under the *HRR25* promoter was analyzed by fluorescence microscopy. **(B)** Cells coexpressing Atg39^167-398^-GFP and the lipid droplet protein Osw5 labeled with mCherry were incubated in oleate-containing medium for 9 h to induce large lipid droplet formation and analyzed under a fluorescence microscope. **(C)** The 4×HA sequence was inserted between the APH^297-324^ and APH^365-379^ of Atg39, and cells expressing this Atg39 variant were analyzed by immunoelectron microscopy using anti-HA antibody. Immunogold signals in the vicinity of the INM and ONM were counted and the results are shown in the graph. Scale bars, 5 µm (A, B), 200 nm (C).

**Figure S2.**
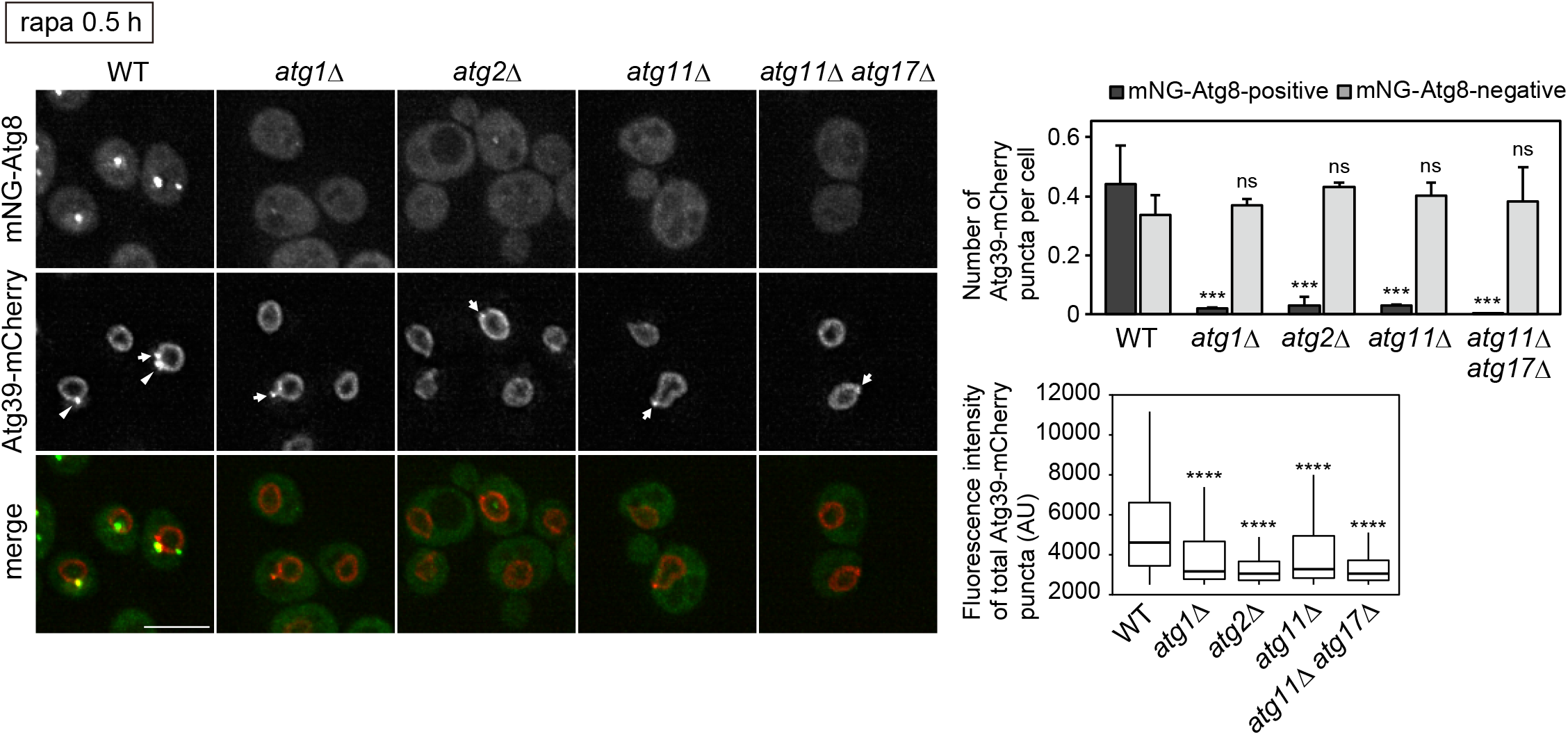
Atg39 assembly in different *atg* mutants. Cells constitutively expressing Atg39-mCherry under the *HRR25* promoter were treated with rapamycin for 0.5 h and analyzed by fluorescence microscopy. The number of Atg39-mCherry puncta per cell (top right) and the fluorescence intensity of Atg39-mCherry puncta (including both mNG-Atg8-positive and -negative puncta) (bottom right) are shown. Arrow heads, mNG-Atg8-positive Atg39 puncta. Arrows, mNG-Atg8-negative Atg39 puncta. Bars represent means ± s.d. (n = 3). \*\*\**P* < 0.001 (Student’s *t*-test) (top right). \*\*\*\**P* < 0.001 (Mann–Whitney *U* test) (bottom right). Scale bars, 5 µm

**Figure S3.**
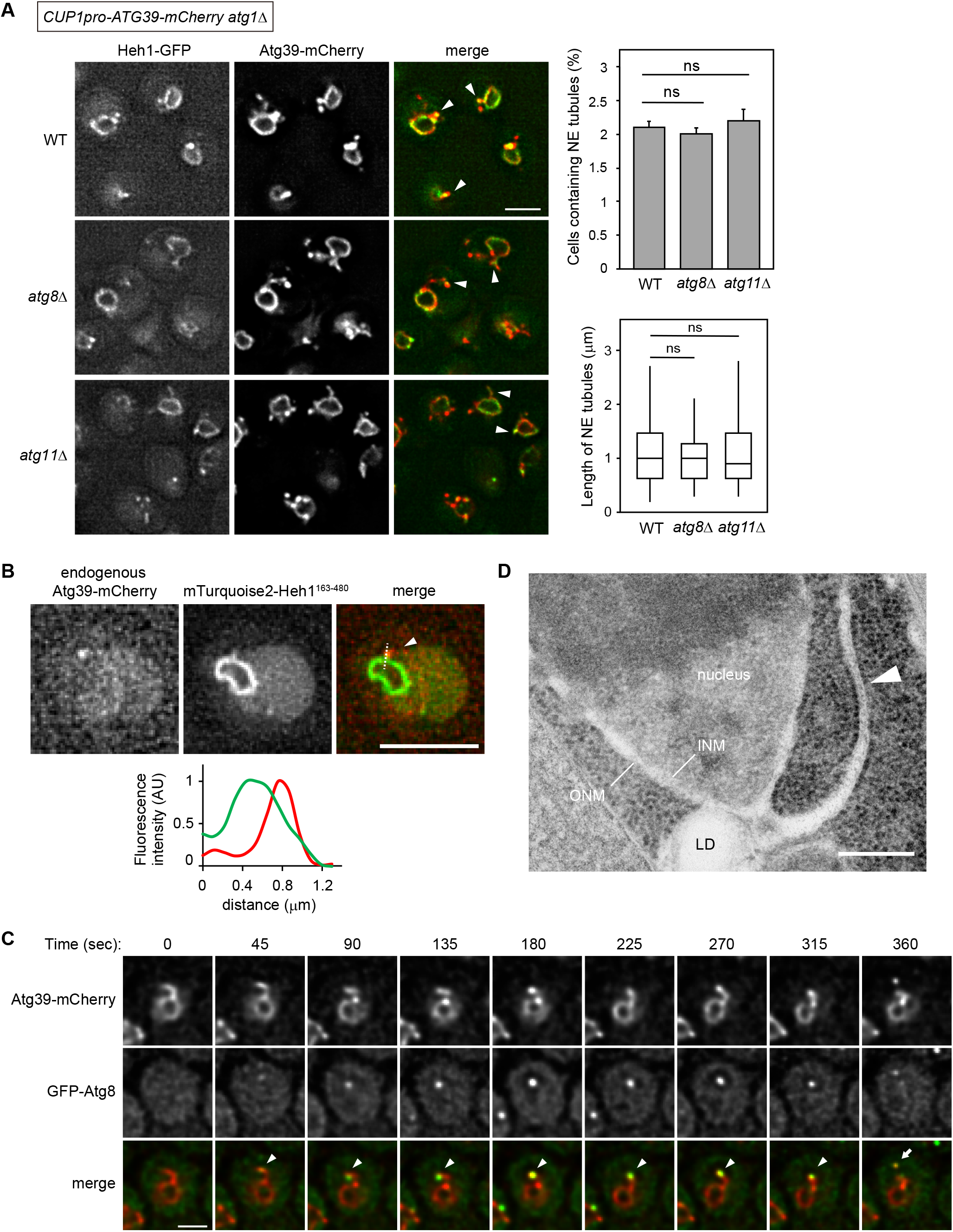
Analysis of NE tubules induced by Atg39 overexpression. **(A)** NE tubule formation was induced by Atg39 overexpression and 2 h rapamycin treatment in *atg1*Δ cells, and the effects of additional deletion of *ATG8* or *ATG11* on tubule formation was examined by fluorescence microscopy. The percentage of cells containing NE tubules (top right) and the length of NE tubules (bottom right) are shown. Bars represent means ± s.d. (n = 3). Statistical significance was determined using Tukey’s multiple comparisons test (top right) and the Kruskal-Wallis test (bottom right), respectively. Arrowheads, NE tubules with GFP signals. **(B)** Cells expressing Atg39-mCherry under the own promoter at the original chromosomal locus were treated with rapamycin for 4 h, and Atg39-mCherry puncta formed at the NE protrusion tip were analyzed by fluorescence microscopy. Fluorescence intensity along the dashed line was measured and is shown graphically. Arrowhead, NE protrusion. **(C)** Cells expressing Atg39-mCherry under the *CUP1* promoter were treated with rapamycin for 20 min and then fluorescence images were taken at 15-s intervals. Arrowheads, GFP-Atg8-positive Atg39-mCherry puncta at the tip of NE tubules. Arrow, NDV. **(D)** A representative EM image of a NE tubule without the INM (arrowhead) generated in *atg1*Δ cells overexpressing Atg39 and treated with rapamycin treatment for 2 h. Scale bars, 5 µm (A, B), 2 µm (C), 200 nm (D)

**Figure S4.**
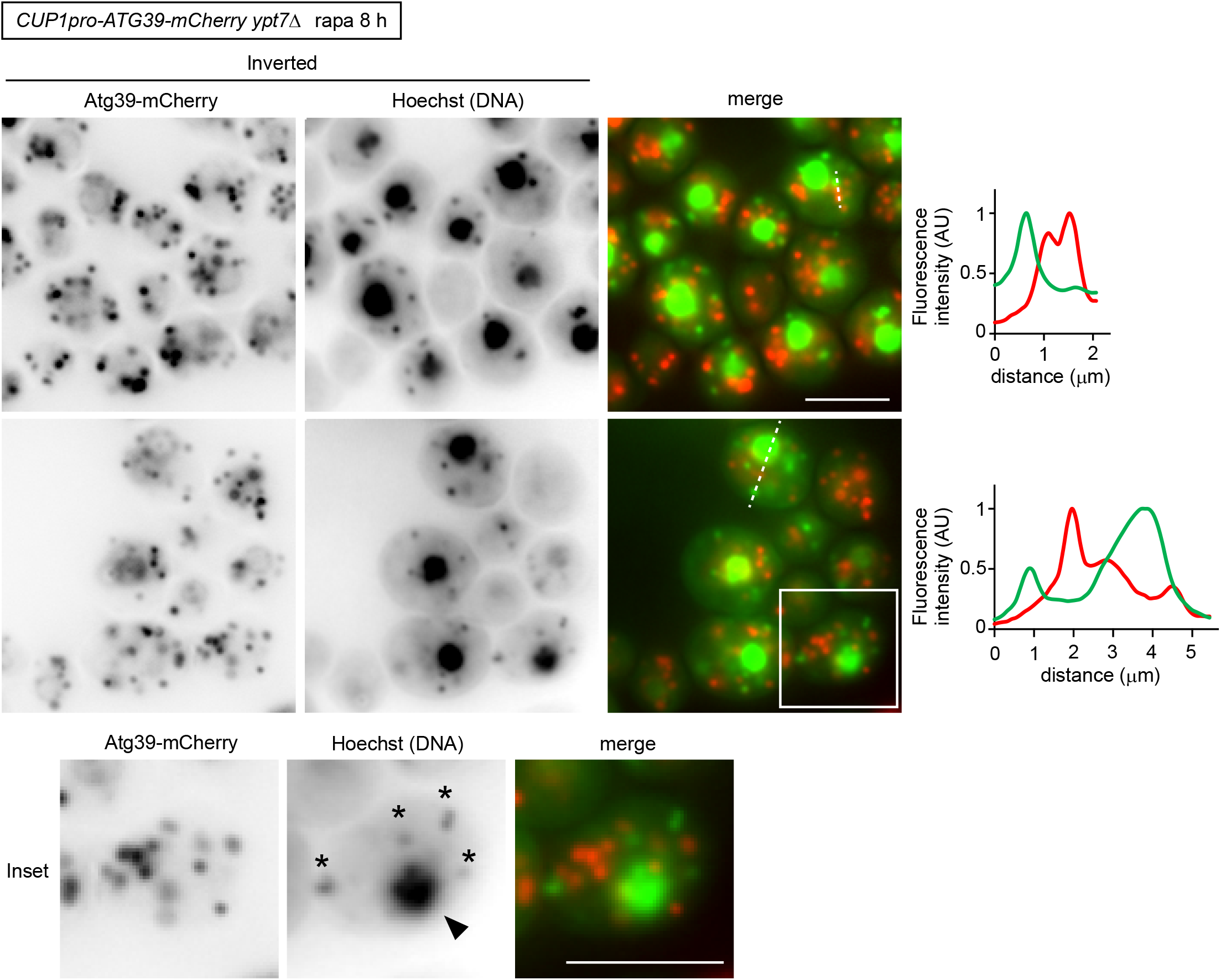
NDVs do not contain DNA. *ypt7*Δ cells overexpressing Atg39-mCherry were treated with rapamycin for 8 h, and the DNA was stained with Hoechst 33258. Arrowheads, nuclei. Small puncta marked by asterisks and that did not colocalize with Atg39-mCherry puncta represent mitochondrial DNA. Fluorescence intensity along the dashed line was measured and is shown in the graph. Scale bars, 5 µm.

**Table S1.** Yeast strains used in this study.

| Name | Genotype |
| --- | --- |
| YKM976 | BJ3505 *natNT2-ADHpro-3HA-ATG39-EGFP-hphNT1 atg1Δ::zeoNT3* |
| YKM977 | BJ3505 *natNT2-ADHpro-ATG39-EGFP-hphNT1 atg1Δ::zeoNT3* |
| YKM1932 | BY4741 *natNT2-HRR25pro-GFP<sup>11</sup>-ATG39* |
| YKM1932 | BY4741 *natNT2-HRR25pro-GFP<sup>11</sup>-ATG39* |
| YKM1938 | BY4741 *HEH1-GFP<sup>11</sup>-natNT2* |
| YKM1536 | BY4741 *TAL1-EGFP-kanMX6 atg39Δ::natNT2* |
| YKM1998 | BY4741 *natNT2-HRR25pro-ATG39-mCherry-hphNT1 his3::mNeonGreen-ATG8-kanMX4* |
| YKM1094 | BY4741 *natNT2-CUP1pro-ATG39-mCherry-hphNT1 his3::GFP-ATG8<sup>G116</sup>-zeoNT3* |
| YKY156 | BY4741 *natNT2-CUP1pro-ATG39<sup>1-296</sup>-mCherry-kanMX6 his3::GFP-ATG8<sup>G116</sup>-zeoNT3* |
| YKM1329 | BY4741 *natNT2-CUP1pro-ATG39-mCherry-hphNT1 HEH1-EGFP-kanMX4 atg1Δ::zeoNT3* |
| YOT61 | BY4741 *atg39Δ::natNT2 atg1Δ::zeoNT3 his3::pRS303-CUP1pro-ATG39-mCherry* |
| YOT62 | BY4741 *atg39Δ::natNT2 atg1Δ::zeoNT3 his3::pRS303-CUP1pro-ATG39<sup>1-296</sup>-mCherry* |
| YOT37 | BY4741 *natNT2-CUP1pro-ATG39-mCherry-hphNT1 TAL1-EGFP-zeoNT3 atg1Δ::URA3* |
| YOT38 | BY4741 *natNT2-CUP1pro-ATG39-mCherry-hphNT1 NOP56-EGFP-zeoNT3 atg1Δ::URA3* |
| YOT22 | BY4741 *natNT2-CUP1pro-ATG39-mCherry-hphNT1 HTA2-EGFP-kanMX6 atg1Δ::URA3* |
| YKM1512 | BY4741 *natNT2-CUP1pro-ATG39-mCherry-hphNT1 TAL1-EGFP-kanMX6 ypt7Δ::zeoNT3* |
| YKM1506 | BY4741 *natNT2-CUP1pro-ATG39-mCherry-hphNT1 HTA2-EGFP-kanMX6 ypt7Δ::zeoNT3* |
| YKM1515 | BY4741 *natNT2-CUP1pro-ATG39-mCherry-hphNT1 HTZ1-EGFP-kanMX6 ypt7Δ::zeoNT3* |
| YKM1809 | BY4741 *natNT2-HRR25pro-ATG39-EGFP-kanMX6 PUS1-mCherry-hphNT1* |
| YKM1813 | BY4741 *natNT2-HRR25pro-ATG39<sup>136-398</sup>-EGFP-kanMX6 PUS1-mCherry-hphNT1* |
| YKM1814 | BY4741 *natNT2-HRR25pro-ATG39<sup>167-398</sup>-EGFP-kanMX6 PUS1-mCherry-hphNT1* |
| YKM1857 | BY4741 *natNT2-HRR25pro-ATG39<sup>167-398</sup>-EGFP-kanMX6 OSW5-mCherry-hphNT1* |
| YKM2040 | BY4741 *natNT2-HRR25pro-ATG39-mCherry-hphNT1 his3::mNeonGreen-ATG8-kanMX4 atg1Δ::zeoNT3* |
| YKM2041 | BY4741 *natNT2-HRR25pro-ATG39-mCherry-hphNT1 his3::mNeonGreen-ATG8-kanMX4 atg2Δ::zeoNT3* |
| YKM2036 | BY4741 *natNT2-HRR25pro-ATG39-mCherry-hphNT1 his3::mNeonGreen-ATG8-kanMX4 atg11Δ::zeoNT3* |
| YKM2043 | BY4741 *natNT2-HRR25pro-ATG39-mCherry-hphNT1 his3::mNeonGreen-ATG8-kanMX4 atg11Δ::zeoNT3 atg17Δ::CgHIS3* |
| YOT43 | BY4741 *natNT2-CUP1pro-ATG39-mCherry-hphNT1 HEH1-EGFP-kanMX4 atg1Δ::URA3 atg8Δ::zeoNT3* |
| YOT44 | BY4741 *natNT2-CUP1pro-ATG39-mCherry-hphNT1 HEH1-EGFP-kanMX4 atg1Δ::URA3 atg11Δ::zeoNT3* |
| YKM1990 | BY4741 *ATG39-mCherry-hphNT his3::pRS303-HRR25pro-mTurquoise2-HEH1<sup>163-480</sup>* |
| YKM1138 | BY4741 *natNT2-CUP1pro-ATG39-mCherry-hphNT1 ypt7Δ::zeoNT3* |

**Table S2.**
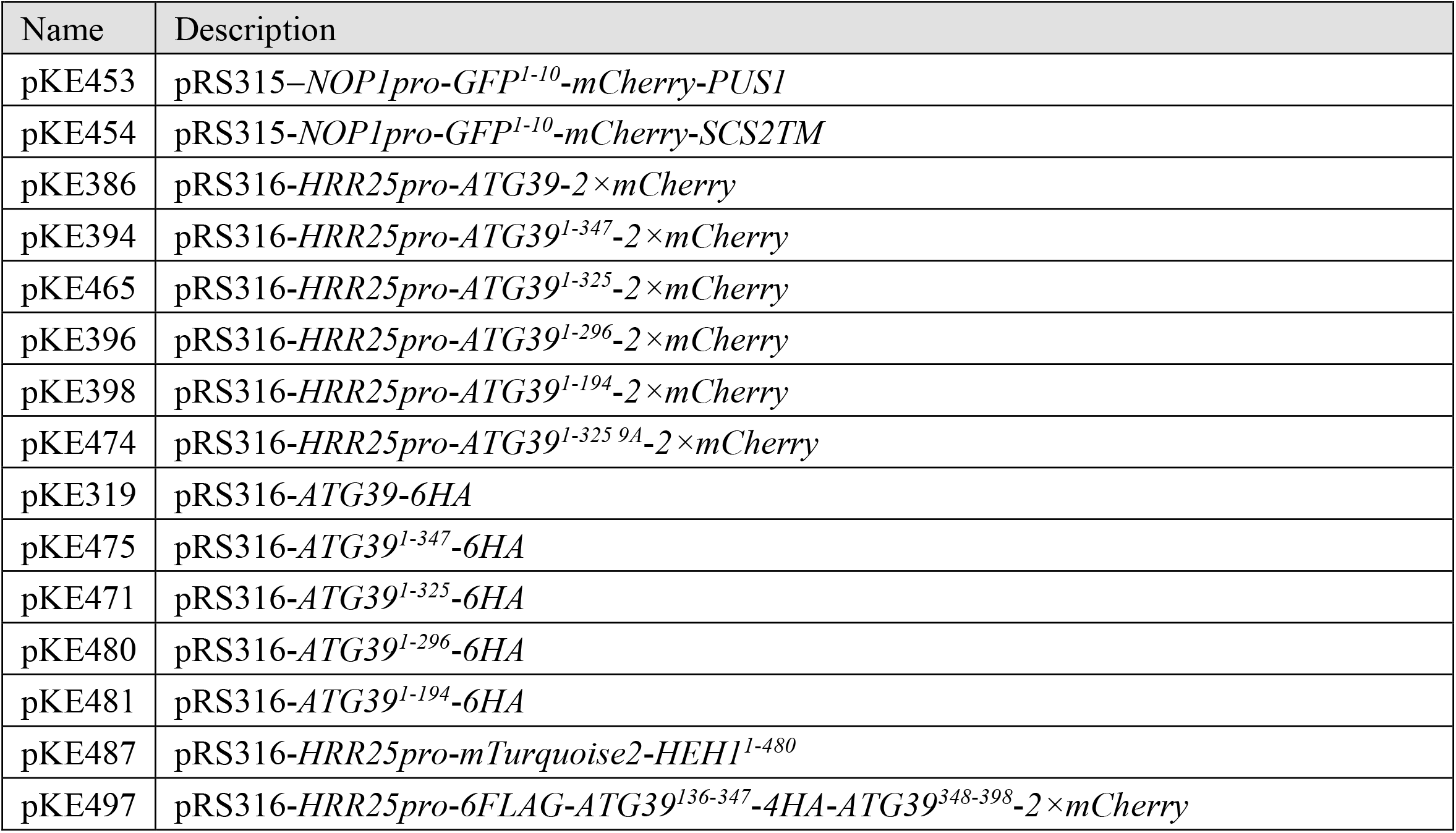
Plasmids used in this study.

## References

Arii, J., M. Watanabe, F. Maeda, N. Tokai-Nishizumi, T. Chihara, M. Miura, Y. Maruzuru, N. Koyanagi, A. Kato, and Y. Kawaguchi. 2018. ESCRT-III mediates budding across the inner nuclear membrane and regulates its integrity. Nat. Commun. 9. doi:10.1038/s41467-018-05889-9.

Baumgart, T., B.R. Capraro, C. Zhu, and S.L. Das. 2011. Thermodynamics and mechanics of membrane curvature generation and sensing by proteins and lipids. Annu. Rev. Phys. Chem. 62:483–506. doi:10.1146/annurev.physchem.012809.103450.

Breker, M., M. Gymrek, O. Moldavski, and M. Schuldiner. 2014. LoQAtE—Localization and Quantitation ATlas of the yeast proteomE. A new tool for multiparametric dissection of single-protein behavior in response to biological perturbations in yeast. Nucleic Acids Res. 42:D726–D730. doi:10.1093/nar/gkt933.

Dou, Z., C. Xu, G. Donahue, T. Shimi, J.-A. Pan, J. Zhu, A. Ivanov, B.C. Capell, A.M. Drake, P.P. Shah, J.M. Catanzaro, M. Daniel Ricketts, T. Lamark, S.A. Adam, R. Marmorstein, W.-X. Zong, T. Johansen, R.D. Goldman, P.D. Adams, and S.L. Berger. 2015. Autophagy mediates degradation of nuclear lamina. Nature. 527:105–109. doi:10.1038/nature15548.

Drin, G., and B. Antonny. 2010. Amphipathic helices and membrane curvature. FEBS Lett. 584:1840–1847. doi:10.1016/j.febslet.2009.10.022.

Enam, C., Y. Geffen, T. Ravid, and R.G. Gardner. 2018. Protein Quality Control Degradation in the Nucleus. Annu. Rev. Biochem. 87:725–749. doi:10.1146/annurev-biochem-062917-012730.

Farré, J.-C., and S. Subramani. 2016. Mechanistic insights into selective autophagy pathways: lessons from yeast. Nat. Rev. Mol. Cell Biol. 17:537–52. doi:10.1038/nrm.2016.74.

Janke, C., M.M. Magiera, N. Rathfelder, C. Taxis, S. Reber, H. Maekawa, A. Moreno-Borchart, G. Doenges, E. Schwob, E. Schiebel, and M. Knop. 2004. A versatile toolbox for PCR-based tagging of yeast genes: New fluorescent proteins, more markers and promoter substitution cassettes. Yeast. doi:10.1002/yea.1142.

Khaminets, A., T. Heinrich, M. Mari, P. Grumati, A.K. Huebner, M. Akutsu, L. Liebmann, A. Stolz, S. Nietzsche, N. Koch, M. Mauthe, I. Katona, B. Qualmann, J. Weis, F. Reggiori, I. Kurth, C.A. Hübner, and I. Dikic. 2015. Regulation of endoplasmic reticulum turnover by selective autophagy. Nature. 522:354–358. doi:10.1038/nature14498.

Kirisako, T., M. Baba, N. Ishihara, K. Miyazawa, M. Ohsumi, T. Yoshimori, T. Noda, and Y. Ohsumi. 1999. Formation Process of Autophagosome Is Traced with Apg8 / Aut7p in Yeast. Jcb. 147:435–446. doi:10.1083/jcb.147.2.435.

Lee, C.P., P.T. Liu, H.N. Kung, M.T. Su, H.H. Chua, Y.H. Chang, C.W. Chang, C.H. Tsai, F.T. Liu, and M.R. Chen. 2012. The ESCRT Machinery Is Recruited by the Viral BFRF1 Protein to the Nucleus-Associated Membrane for the Maturation of Epstein-Barr Virus. PLoS Pathog. 8. doi:10.1371/journal.ppat.1002904.

Lee, C.W., F. Wilfling, P. Ronchi, M. Allegretti, S. Mosalaganti, S. Jentsch, M. Beck, and B. Pfander. 2020. Selective autophagy degrades nuclear pore complexes. Nat. Cell Biol. doi:10.1038/s41556-019-0459-2.

Mao, K., K. Wang, X. Liu, and D.J. Klionsky. 2013. The scaffold protein Atg11 recruits fission machinery to drive selective mitochondria degradation by autophagy. Dev. Cell. 26:9–18. doi:10.1016/j.devcel.2013.05.024.

McCullough, J., A. Frost, and W.I. Sundquist. 2018. Structures, Functions, and Dynamics of ESCRT-III/Vps4 Membrane Remodeling and Fission Complexes. Annu. Rev. Cell Dev. Biol. 34:85–109. doi:10.1146/annurev-cellbio-100616-060600.

Mijaljica, D., and R.J. Devenish. 2013. Nucleophagy at a glance. J. Cell Sci. 126:4325–4330. doi:10.1242/jcs.133090.

Mochida, K., Y. Oikawa, Y. Kimura, H. Kirisako, H. Hirano, Y. Ohsumi, and H. Nakatogawa. 2015. Receptor-mediated selective autophagy degrades the endoplasmic reticulum and the nucleus. Nature. 522:359–362. doi:10.1038/nature14506.

Mochida, K., A. Yamasaki, K. Matoba, H. Kirisako, N.N. Noda, and H. Nakatogawa. 2020. Super-assembly of ER-phagy receptor Atg40 induces local ER remodeling at contacts with forming autophagosomal membranes. Nat. Commun. 11. doi:10.1038/s41467-020-17163-y.

Morishita, H., and N. Mizushima. 2019. Diverse Cellular Roles of Autophagy. Annu. Rev. Cell Dev. Biol. 35:453–475. doi:10.1146/annurev-cellbio-100818-125300.

Nakatogawa, H. 2020. Mechanisms governing autophagosome biogenesis. Nat. Rev. Mol. Cell Biol. doi:10.1038/s41580-020-0241-0.

Schindelin, J., I. Arganda-Carreras, E. Frise, V. Kaynig, M. Longair, T. Pietzsch, S. Preibisch, C. Rueden, S. Saalfeld, B. Schmid, J.Y. Tinevez, D.J. White, V. Hartenstein, K. Eliceiri, P. Tomancak, and A. Cardona. 2012. Fiji: An open-source platform for biological-image analysis. Nat. Methods. 9:676–682. doi:10.1038/nmeth.2019.

Sikorski, R.S., and P. Hieter. 1989. A system of shuttle vectors and yeast host strains designed for efficient manipulation of DNA in Saccharomyces cerevisiae. Genetics. doi:0378111995000377 [pii].

Smoyer, C.J., S.S. Katta, J.M. Gardner, L. Stoltz, S. McCroskey, W.D. Bradford, M. McClain, S.E. Smith, B.D. Slaughter, J.R. Unruh, and S.L. Jaspersen. 2016. Analysis of membrane proteins localizing to the inner nuclear envelope in living cells. J. Cell Biol. 215:575–590. doi:10.1083/jcb.201607043.

Stachowiak, J.C., E.M. Schmid, C.J. Ryan, H.S. Ann, D.Y. Sasaki, M.B. Sherman, P.L. Geissler, D.A. Fletcher, and C.C. Hayden. 2012. Membrane bending by protein-protein crowding. Nat. Cell Biol. 14:944–9. doi:10.1038/ncb2561.

Tomioka, Y., T. Kotani, H. Kirisako, Y. Oikawa, Y. Kimura, H. Hirano, Y. Ohsumi, and H. Nakatogawa. 2020. TORC1 inactivation stimulates autophagy of nucleoporin and nuclear pore complexes. J. Cell Biol. 219. doi:10.1083/jcb.201910063.

Vevea, J.D., E.J. Garcia, R.B. Chan, G. Di Paolo, J.M. Mccaffery, L.A. Pon, J.D. Vevea, E.J. Garcia, R.B. Chan, B. Zhou, M. Schultz, G. Di Paolo, J.M. Mccaffery, and L.A. Pon. 2015. Role for Lipid Droplet Biogenesis and Microlipophagy in Adaptation to Lipid Imbalance in Article Role for Lipid Droplet Biogenesis and Microlipophagy in Adaptation to Lipid Imbalance in Yeast. Dev. Cell. 35:584–599. doi:10.1016/j.devcel.2015.11.010.

Webster, B.M., P. Colombi, J. Jäger, and C. Patrick Lusk. 2014. Surveillance of nuclear pore complex assembly by ESCRT-III/Vps4. Cell. doi:10.1016/j.cell.2014.09.012.

West, M., N. Zurek, A. Hoenger, and G.K. Voeltz. 2011. A 3D analysis of yeast ER structure reveals how ER domains are organized by membrane curvature. J. Cell Biol. 193:333–346. doi:10.1083/jcb.201011039.

Woulfe, J. 2008. Nuclear bodies in neurodegenerative disease. Biochim. Biophys. Acta. 1783:2195–206. doi:10.1016/j.bbamcr.2008.05.005.

